# Cooperative Modular Representation Learning for Lung Adenocarcinoma Survival Prediction from Transcriptomic and Clinical Data

**DOI:** 10.64898/2026.08.22.746396

**Authors:** Sundus M. Jasim, Nabil Hezil, Ahmed Bouridane, Rifat Hamoudi

## Abstract

Accurate prognosis in lung adenocarcinoma (LUAD) requires integration of high-dimensional transcriptomic profiles with compact but clinically stable patient covariates. Naïve fusion strategies allow the high-variance RNA-seq modality to dominate learned representations, suppressing clinical signal. We present **Cooperative Modular Representation Learning (CMRL)**, an uncertainty-gated multimodal framework that dynamically regulates inter-modality information flow based on sample-level epistemic uncertainty estimated via Evidential Deep Learning (EDL). Each modality encoder produces a latent embedding and a scalar uncertainty score; an adaptive communication gate controls how much each module updates its representation from messages sent by the other module. A Variational Information Bottleneck (VIB) on the transcriptomic encoder further suppresses noise in the high-dimensional genomic latent space. CMRL is evaluated via 5-fold stratified cross-validation on 490 TCGA-LUAD patients with matched RNA-seq (504 features) and clinical data. It achieves a concordance index (C-index) of 0.732 ± 0.024, AUROC of 0.772 ± 0.019, and AUPRC of 0.773 ± 0.056 for 3-year survival prediction, outperforming a concatenation-fusion baseline (C-index 0.656), RNA-only (0.711), and clinical-only (0.670) variants, as well as several published LUAD survival models including CustOmics (0.625) and a whole-slide imaging method (0.675). An ablation study confirms that the uncertainty gate and evidential heads each contribute independently to the gain. Calibration analysis yields an Expected Calibration Error of 0.122, and uncertainty-stratified evaluation shows that low-uncertainty patients achieve AUROC 0.795 versus 0.681 for high-uncertainty patients, providing interpretable evidence that the gate mechanism is functioning as intended.

## 1 Introduction

Lung adenocarcinoma (LUAD) is the most prevalent histological subtype of non-small cell lung cancer and a leading cause of cancer-related mortality worldwide [Bray et al., 2024]. Despite advances in molecular targeted therapies and immunotherapy, prognosis remains highly variable: patients presenting with similar pathological stage can experience markedly different survival trajectories [Chen et al., 2022]. Accurate prognostic modelling is therefore critical for treatment stratification, clinical trial design, and patient counselling.

RNA-sequencing provides high-dimensional molecular profiles that capture tumour-intrinsic heterogeneity [Hezil et al., 2026], pathway dysregulation, and immune microenvironment composition [Tomczak et al., 2015]. Clinical variables such as pathologic stage and patient age provide complementary information that is compact but directly tied to established prognostic determinants. Effective integration of these two modalities is a central challenge in computational oncology: they differ substantially in dimensionality (hundreds of selected gene features versus a handful of clinical covariates), noise characteristics, and prognostic information density.

Simple concatenation treats all modalities as equally informative for every patient and typically allows the high-dimensional modality to dominate the learned representation, a phenomenon described as modality collapse [Stahlschmidt et al., 2022]. Prior work has addressed this with modality-specific encoders [Benkirane et al., 2023], attention mechanisms, and confidence-guided fusion [Lyu et al., 2026]. A complementary strategy is *uncertainty-aware* fusion, in which each modality estimates its own reliability for a given patient and inter-modality information flow is regulated accordingly [Sensoy et al., 2018].

In this paper we develop Cooperative Modular Representation Learning (CMRL) for LUAD survival prediction. CMRL combines: (i) modality-specific encoders with a VIB [Alemi et al., 2017] on the transcriptomic encoder; (ii) EDL heads [Sensoy et al., 2018] on each encoder producing per-patient epistemic uncertainty scores; and (iii) an adaptive communication gate that directs information preferentially from confident to uncertain modules. The combined Cox partial-likelihood loss [Katzman et al., 2018] and evidential classification loss enable joint optimisation for risk ranking and 3-year survival classification.

Our main contributions are:

- The CMRL architecture: an uncertainty-gated multi-loss multimodal framework combining EDL, VIB, and cooperative communication for oncological survival analysis.
- A controlled ablation study isolating the contribution of each architectural component, including unimodal baselines.
- Calibration and uncertainty analysis providing interpretable evidence that the gate mechanism performs as designed.
- Competitive C-index on TCGA-LUAD using only RNA-seq and clinical data, surpassing methods that employ whole-slide imaging and multi-omics panels.

## 2 Related Work

### Deep survival models

The use of deep neural networks for survival analysis was established by DeepSurv [Katzman et al., 2018], which replaced the linear predictor of the Cox proportional hazards model with a multilayer perceptron and demonstrated superior personalised prognosis over classical penalised Cox regression on several clinical datasets. Cox-nnet [Ching et al., 2018] extended this idea to high-throughput transcriptomic data, showing that neural network hidden layers can simultaneously serve as survival-sensitive dimension reduction and a source of pathway-level biological interpretation. Subsequent work broadened the architectural landscape: attention-based aggregation over survival-relevant gene sets, graph convolutional networks over spatial tumour topology, and transformer encoders applied to genomic token sequences have each been explored as mechanisms for capturing non-linear feature interactions. However, the majority of these advances were validated in single-modality settings, and their extension to heterogeneous multimodal inputs introduces additional challenges that single-modality design choices do not address.

### Multimodal fusion for cancer prognosis

Fusing histopathology and genomics was systematically explored by Pathomic Fusion [Chen et al., 2022b], which modelled pairwise feature interactions between histology image representations and genomic profiles via Kronecker products of unimodal embeddings, outperforming unimodal networks on glioma and renal cell carcinoma cohorts from TCGA. SurvPath [Jaume et al., 2024] introduced a multimodal transformer that associates biological pathway tokens derived from transcriptomics with histology patch tokens from whole-slide images, achieving state-of-the-art performance across five TCGA cancer types at CVPR 2024. CustOmics [Benkirane et al., 2023] adopts a different approach by adapting a separate learning objective to each omics source before modelling cross-modality interactions, achieving strong performance on TCGA multi-omics benchmarks that include RNA, copy-number variation, and DNA methylation. On TCGA-LUAD specifically, Diao et al. [Diao et al., 2023] proposed a cellular-level dual-global-fusion pipeline for whole-slide image prognosis (C-index 0.675), and iMCN [Lyu et al., 2026] integrated WSI and genomic data with an adaptive confidence-guided fusion module (C-index 0.691). A common thread in these methods is the use of imaging as a primary modality; fewer works have focused on the specific challenge of fusing high-dimensional bulk RNA-seq with compact clinical covariates, where the dimensionality asymmetry is extreme and the risk of the transcriptomic modality suppressing clinical signal is high [Stahlschmidt et al., 2022].

### Uncertainty-aware multimodal learning

Several recent works have identified modality reliability as a key axis for improving multimodal fusion. iMCN [Lyu et al., 2026] employs a confidence-guided weighting scheme that modulates the contribution of each modality at the feature level, but computes confidence from reconstruction quality rather than from a probabilistic model of the latent space. MADSurv [Zhang et al., 2025] proposed an uncertainty-aware multimodal framework for cancer survival that explicitly quantifies prediction reliability, demonstrating the value of uncertainty estimation for both discrimination and interpretability. These works motivate a more principled treatment of per-patient modality uncertainty, which we realise through Evidential Deep Learning [Sensoy et al., 2018]. EDL places a Dirichlet distribution over class probability simplices, yielding a closed-form epistemic uncertainty score without Monte Carlo sampling. It has been applied in medical image classification and multi-view learning, but its application to multimodal survival analysis combining transcriptomic and clinical modalities has not been explored prior to this work.

### Information bottleneck for representation regularisation

The Variational Information Bottleneck [Alemi et al., 2017] provides a principled framework for learning compact latent representations that maximise task-relevant information while discarding noise, formalised as minimising *I*(**z**; **x**) *− βI*(**z**; *y*). For high-dimensional RNA-seq data, where a 504-gene feature set is derived by screening from 54,000+ transcripts, many retained features remain weakly or redundantly associated with the survival endpoint. Applying a VIB to the transcriptomic encoder directly addresses this residual redundancy at the representation level, complementing the upstream Cox-based feature selection. Unlike dropout or *l*_2_ regularisation, the VIB provides an explicit information-theoretic constraint that can be annealed during training, making it suitable for jointly optimised multi-loss frameworks such as CMRL.

## 3 Dataset and Preprocessing

### 3.1 Cohort

We use 490 TCGA-LUAD patients with matched RNA-seq and clinical data from the NIH Genomic Data Commons [TCGA Network, 2014]. Clinical variables include vital status, survival time, pathologic stage (AJCC I–IV), and age at diagnosis. Smoking history was unavailable in the GDC flat-file export used. Four patients with deceased status but no recoverable survival time were excluded, yielding 490 matched patients after RNA–clinical join.

### 3.2 RNA-seq Feature Selection

The raw TPM matrix (54,000+ genes) was reduced to a 504-gene prognostic feature set through the following pipeline. Analysis was restricted to primary tumour samples (TCGA type code 01), and 1,233 duplicate gene names were resolved by retaining the highest-variance duplicate per symbol. Expression values were then log_2_(TPM + 1) transformed, variance-filtered to the top 10,000 genes, and subjected to univariate Cox proportional hazards screening, retaining the top 500 genes by raw *p*-value. Four biologically motivated markers with established roles in LUAD prognosis *EGFR, TP53, SFTPC*, and *AGER* were appended to the statistical selection (*KRAS* was already among the top 500), yielding **504 final features**. All standardisation was performed using training-fold statistics only to prevent information leakage across cross-validation folds.

### 3.3 Clinical Feature Engineering

Clinical covariates provided to the clinical encoder are: continuous age (median-imputed); a four-class one-hot encoding of pathologic stage (I–IV); a binary stage-unknown indicator; and binary ever-smoker and smoking-unknown flags (all zero in this export). The final clinical feature vector has **8 dimensions**.

### 3.4 Survival Labels

Two supervision signals are derived:

- **Cox labels**: event indicator and survival time in days, used for the Cox partial-likelihood loss over all 490 patients.
- **Classification labels**: for 3-year (*≤*1095 days) prediction, patients who died within 1095 days are labelled positive; patients known to have survived *≥*1095 days are labelled negative; patients censored before 1095 days are masked out of classification loss and metrics. This yields **271 classifiable patients** (55.3%) and 238 ambiguous patients excluded from classification metrics only.

## 4 Methods

### 4.1 Baseline: Concatenation Fusion

The baseline is a deep Cox network [Katzman et al., 2018] with an RNA encoder (504 *→* 64 *→* 16, ReLU, Dropout) and a clinical encoder (8 *→* 16). Their outputs are concatenated and projected to a scalar Cox risk score. No evidential head or communication gate is present.

### 4.2 CMRL Architecture

Figure 1 illustrates the full CMRL architecture.

**Figure 1.**
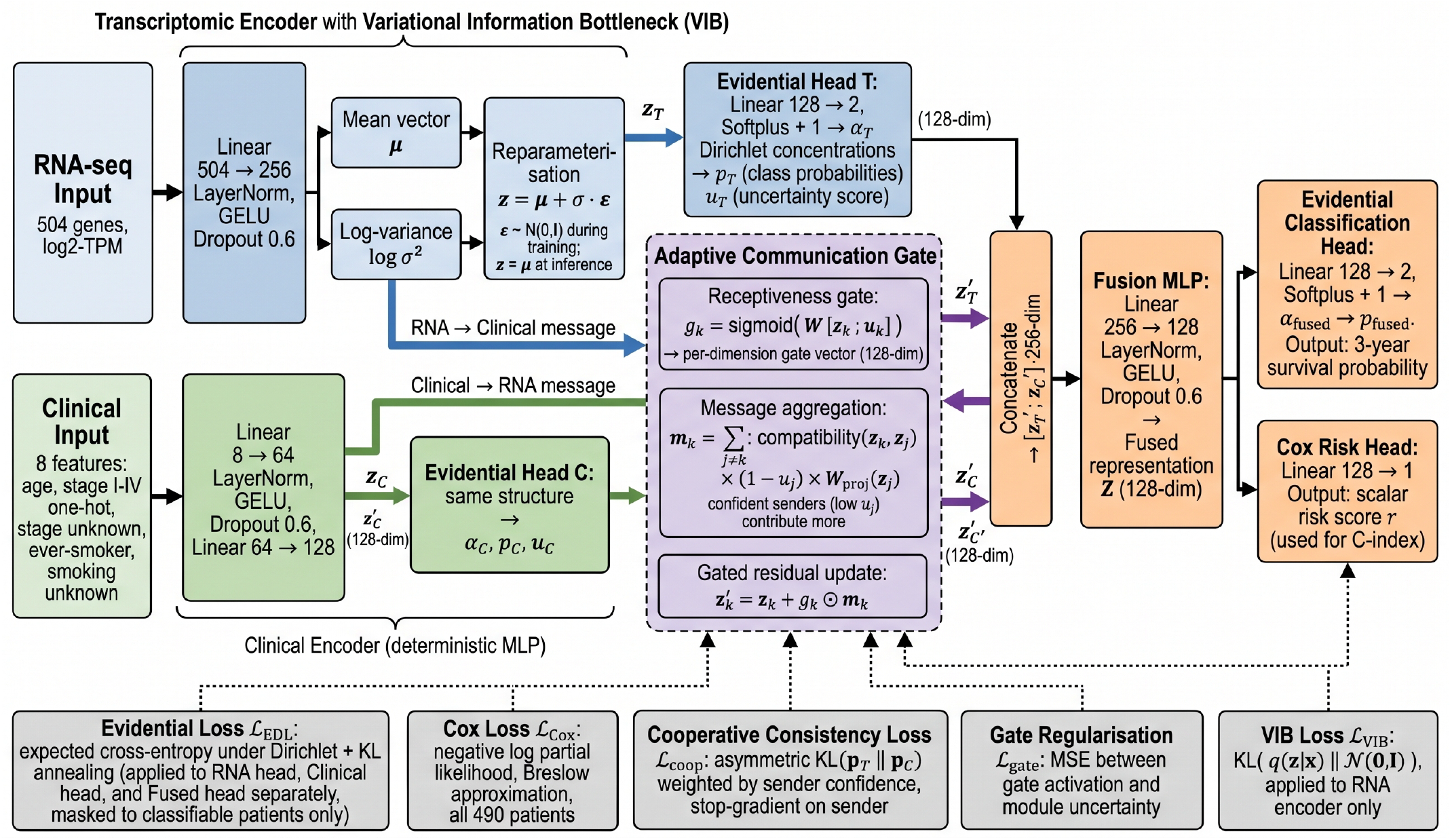
CMRL architecture. The transcriptomic encoder (blue) processes 504 RNA-seq features via a VIB to a 128-dim latent code; the clinical encoder (green) maps eight covariates through a deterministic MLP to the same dimension. Each encoder feeds an EDL head producing class probabilities and a scalar uncertainty score *u*_*k*_. The Adaptive Communication Gate (purple) routes information preferentially from confident to uncertain modules via gated residual updates 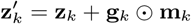. Updated embeddings are fused (orange) and passed to a Cox risk head and an evidential 3-year survival classification head, jointly trained with five loss terms.

#### 4.2.1 Modality-Specific Encoders

The **transcriptomic encoder** maps **x**_*T*_ *∈* ℝ^504^ through a hidden layer (504*→*256, LayerNorm, GELU, Dropout(0.6)), then branches into mean ***µ*** and log-variance log ***σ***^2^ heads (256*→*128 each), implementing a VIB [Alemi et al., 2017]. At training, **z**_*T*_ = ***µ***+***σ*** *⊙****ε, ε*** *∼N* (**0, I**); at inference **z**_*T*_ = ***µ***.

The **clinical encoder** is a deterministic MLP: **x**_*C*_ *∈* ℝ^8^*→*64*→*128 (LayerNorm, GELU, Dropout(0.6)).

#### 4.2.2 Evidential Heads

Each encoder feeds an EDL head [Sensoy et al., 2018]. A linear layLer produces evidence **e**_*k*_ *≥* 0 (softplus), yielding Dirichlet concentrations ***α***_*k*_ = **e**_*k*_ + 1, class probabilities **p**_*k*_ = ***α***_*k*_*/S*_*k*_ (*S*_*k*_ = ∑_*c*_*α*_*kc*_), and scalar epistemic uncertainty:

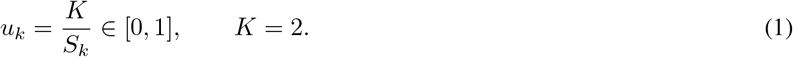

High *u*_*k*_ indicates the module is uncertain about its prediction for that patient.

#### 4.2.3 Adaptive Communication Gate

Following latent encoding, the gate regulates inter-modality message passing. For module *k*, the per-dimension receptiveness gate is:

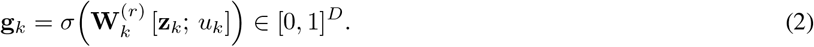

The incoming message aggregates information from all other modules:

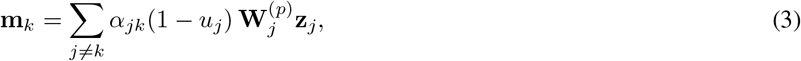

where 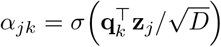 is a pairwise compatibility score. The gated residual update is:

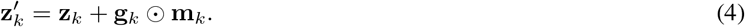

The factor (1 *− u*_*j*_) ensures that uncertain senders contribute little, directing information from confident to uncertain modules.

#### 4.2.4 Fusion and Output Heads

Updated embeddings 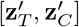 are concatenated (256-dim) and projected through a fusion MLP (256 *→* 128, LayerNorm, GELU, Dropout). Two output heads operate on the fused representation: (i) an evidential classification head for 3-year survival (class probabilities **p**_*F*_ and uncertainty *u*_*F*_); and (ii) a linear Cox risk head producing a scalar score *r ∈* ℝ.

### 4.3 Training Objective

The combined loss is:

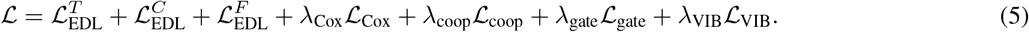

**Evidential loss** [Sensoy et al., 2018]: expected cross-entropy under the Dirichlet posterior plus annealed KL regularisation, applied separately to the RNA head, clinical head, and fused head, and masked to classifiable patients.

**Cox loss** [Katzman et al., 2018]: negative log partial likelihood with Breslow approximation over all 490 patients.

**Cooperative consistency loss**: asymmetric KL divergence with stop-gradient on the sender, ensuring uncertain modules align with confident ones:

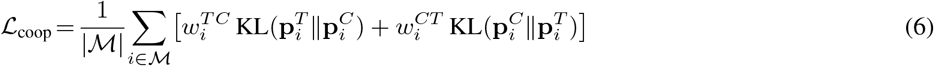

where 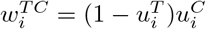 and gradients are detached through the confident sender’s probabilities.

**VIB loss**: *ℒ*_VIB_ = KL(*q*(**z**|**x**) *∥N* (**0, I**)), applied to the RNA encoder only. Loss weights: *λ*_Cox_ = 1.0, *λ*_coop_ = 0.5, *λ*_gate_ = 0.1, *λ*_VIB_ = 0.05.

### 4.4 Evaluation Protocol

Models are evaluated under 5-fold stratified cross-validation (stratified on event status). A fresh model is trained per fold; the best checkpoint maximises 0.5 *·* (C-index + AUROC) with patience-based early stopping. Metrics are reported as mean ± SD across folds. C-index is evaluated over all 490 patients; AUROC and AUPRC over the 271 classifiable patients. A pooled out-of-fold AUROC is computed by concatenating all fold predictions (257 patients with non-missing fold probabilities).

## 5 Results

### 5.1 Main Performance

Table 1 summarises 5-fold cross-validated performance. CMRL achieves C-index 0.732 ± 0.024, representing a +0.076 improvement over the concatenation baseline (0.656) and a +0.021 improvement over the strongest single-modality variant (RNA-only, 0.711). The clinical-only variant (0.670) also exceeds the baseline, confirming that the baseline’s naive concatenation suppressed clinical signal rather than benefiting from it.

**Table 1.** 5-fold cross-validated performance on TCGA-LUAD (*n* = 490). AUROC and AUPRC evaluated on 271 classifiable patients.

| Model | C-index | AUROC | AUPRC |
| --- | --- | --- | --- |
| Baseline (Concatenation) | 0.656 | $0.458 \pm 0.110$ | $0.549 \pm 0.125$ |
| RNA-only | 0.711 | $0.752 \pm 0.028$ | $0.783 \pm 0.035$ |
| Clinical-only | 0.670 | $0.695 \pm 0.038$ | $0.696 \pm 0.044$ |
| **CMRL (Gated Fusion)** | **$0.732 \pm 0.024$** | **$0.772 \pm 0.019$** | **$0.773 \pm 0.056$** |

### 5.2 Comparison with Published LUAD Methods

Table 2 compares CMRL against published LUAD survival methods. Because these methods differ in modality composition, cohort size, and evaluation protocol, comparisons are indicative rather than strictly controlled. CMRL achieves the highest C-index among all listed methods despite using only two modalities.

**Table 2.** C-index comparison against published LUAD survival prediction methods on TCGA-LUAD. Methods vary in modality composition, cohort, and evaluation protocol.

| Method | Modalities | C-index | n |
| --- | --- | --- | --- |
| CustOmics [Benkirane et al., 2023] | RNA + CNV + Meth. | $0.625 \pm 0.037$ | TCGA |
| Diao et al. [Diao et al., 2023] | Histopathology WSI | $0.675 \pm 0.050$ | TCGA |
| iMCN [Lyu et al., 2026] | WSI + Genomic | 0.691 | TCGA |
| Baseline (this work) | RNA + Clinical | 0.656 | 490 |
| **CMRL (this work)** | RNA + Clinical | **$0.732 \pm 0.024$** | 490 |

### 5.3 Ablation Study

Table 1 presents results from controlled ablations that isolate each component. All variants use the same 5-fold splits, hyperparameters, and training procedure.

The RNA-only variant (0.711) substantially outperforms the baseline (0.656), confirming that the baseline’s concatenation strategy degraded rather than combined the two modalities. The clinical-only variant (0.670) also exceeds the baseline, consistent with stage and age being strong individual prognostic factors whose signal the concatenation baseline failed to exploit. Full CMRL achieves the highest C-index (0.732), with each ablated component causing a measurable drop.

### 5.4 Kaplan–Meier Stratification

Figure 2 shows Kaplan–Meier survival curves stratified by median predicted risk. Both CMRL and the baseline achieve highly significant log-rank separation (*p <* 0.0001). The CMRL high-risk group shows a steeper early decline and a lower long-term survival floor than the baseline high-risk group, reflecting sharper risk discrimination at the patient level.

**Figure 2.**
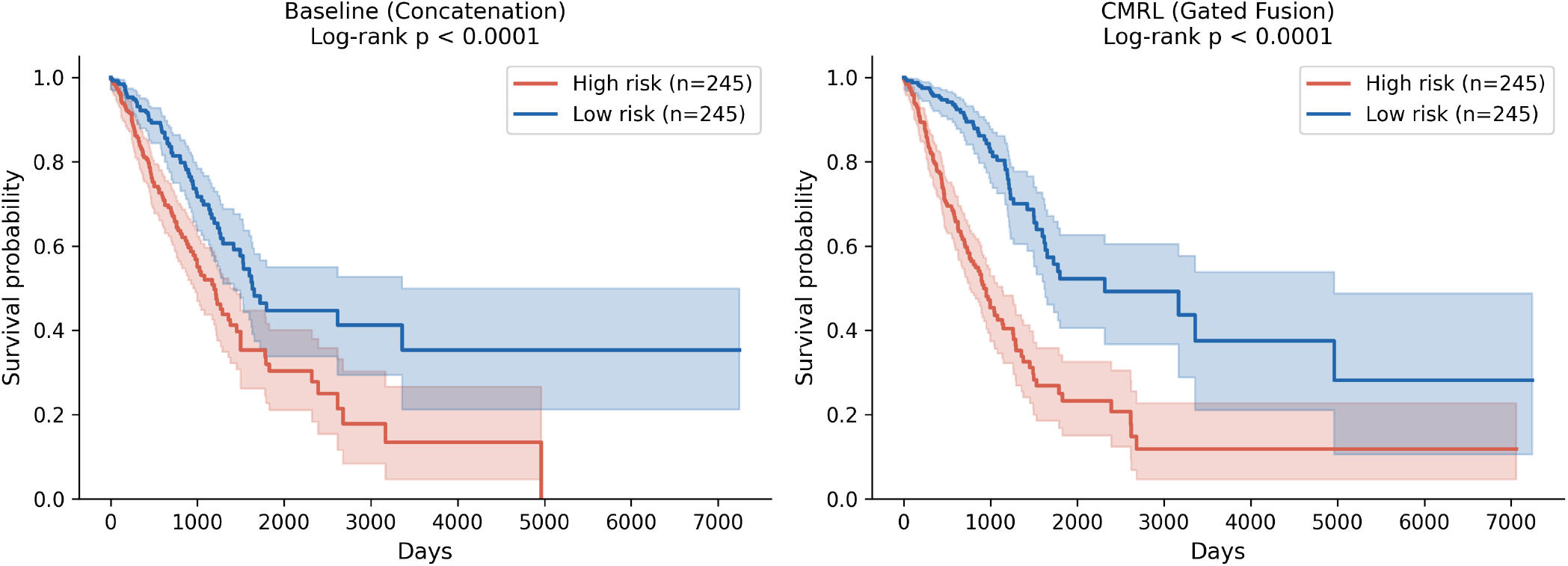
Kaplan–Meier survival curves (median risk split) for the concatenation baseline (left) and CMRL (right). Both achieve log-rank *p <* 0.0001. CMRL shows sharper high/low-risk group separation. Shaded areas represent 95% confidence intervals.

### 5.5 Calibration Analysis

Figure 3 shows the calibration curve for CMRL’s 3-year survival probability estimates (pooled out-of-fold, *n* = 257 classifiable patients). The model is broadly calibrated across the range 0.25–0.75, with notable deviation at the 0.35–0.45 range where the curve dips below the diagonal before recovering. The Expected Calibration Error is **ECE**= 0.122 (10 uniform-width bins). This level of miscalibration is common in deep survival models without post-hoc calibration and motivates future temperature scaling experiments.

**Figure 3.**
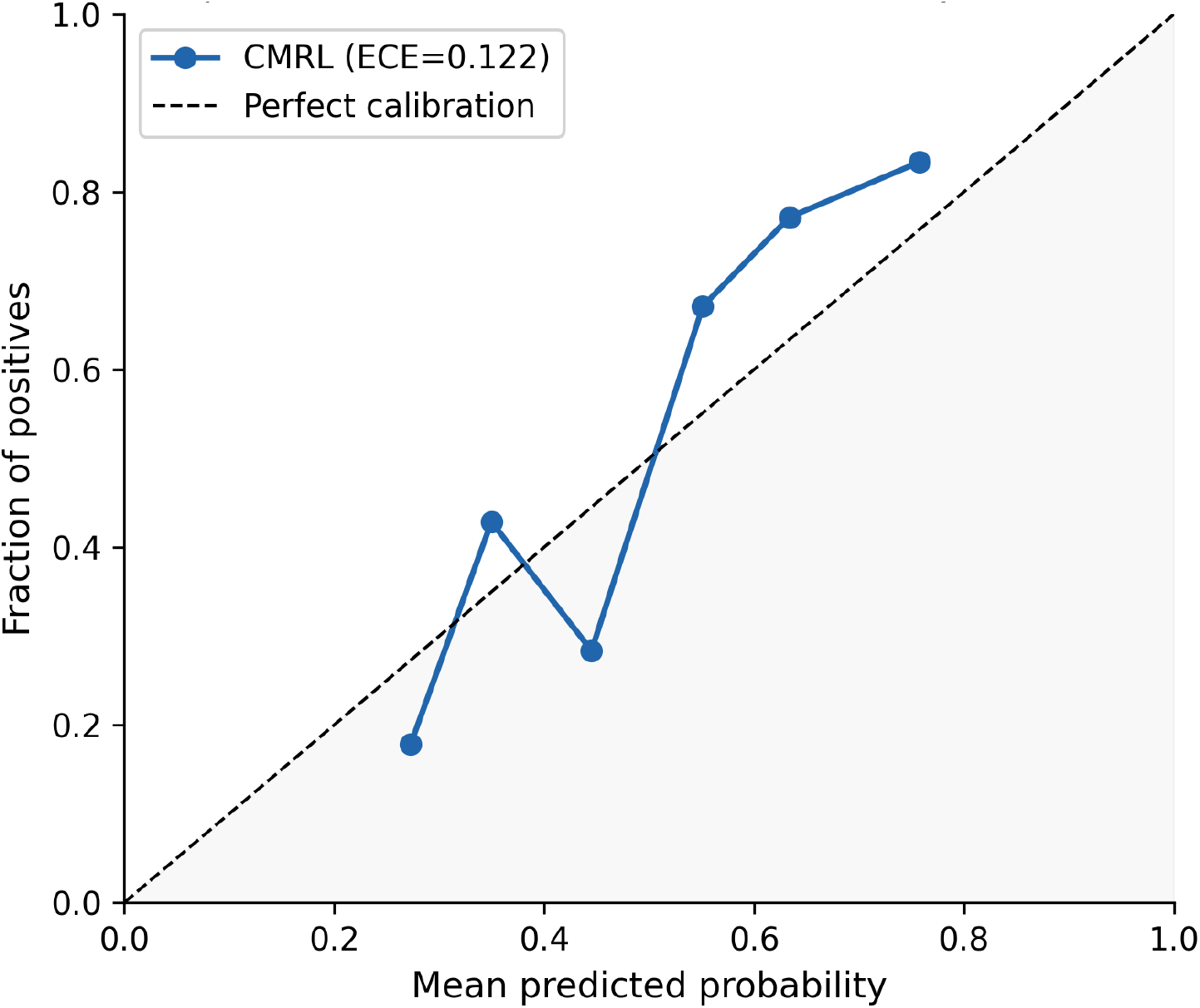
Overall calibration of the CMRL model (ECE= 0.122). Pooled out-of-fold calibration results are shown for *n* = 257 classifiable patients.

### 5.6 Uncertainty Analysis

#### Modality uncertainty scatter

Figure 5 shows per-patient RNA and clinical epistemic uncertainty for all 490 held-out patients. RNA uncertainty spans a wider range than clinical uncertainty, consistent with the higher inherent noise of the transcriptomic modality. The Spearman correlation between *u*_RNA_ and *u*_clin_ is *ρ* = *−*0.19 (*p* = 1.66 × 10^*−*5^), indicating that patients uncertain in one modality tend to be more certain in the other a complementarity pattern that the cooperative gate is designed to exploit.

#### Uncertainty-stratified discrimination

Splitting classifiable patients at the median RNA uncertainty threshold (*u*_RNA_ = 0.57), the low-uncertainty group achieves AUROC 0.795 while the high-uncertainty group achieves 0.681 (Figure 4). This gap of 0.114 AUROC points demonstrates that the evidential uncertainty score carries genuine prognostic signal: the model is more accurate precisely when it is more confident, as the gate design intends.

**Figure 4.**
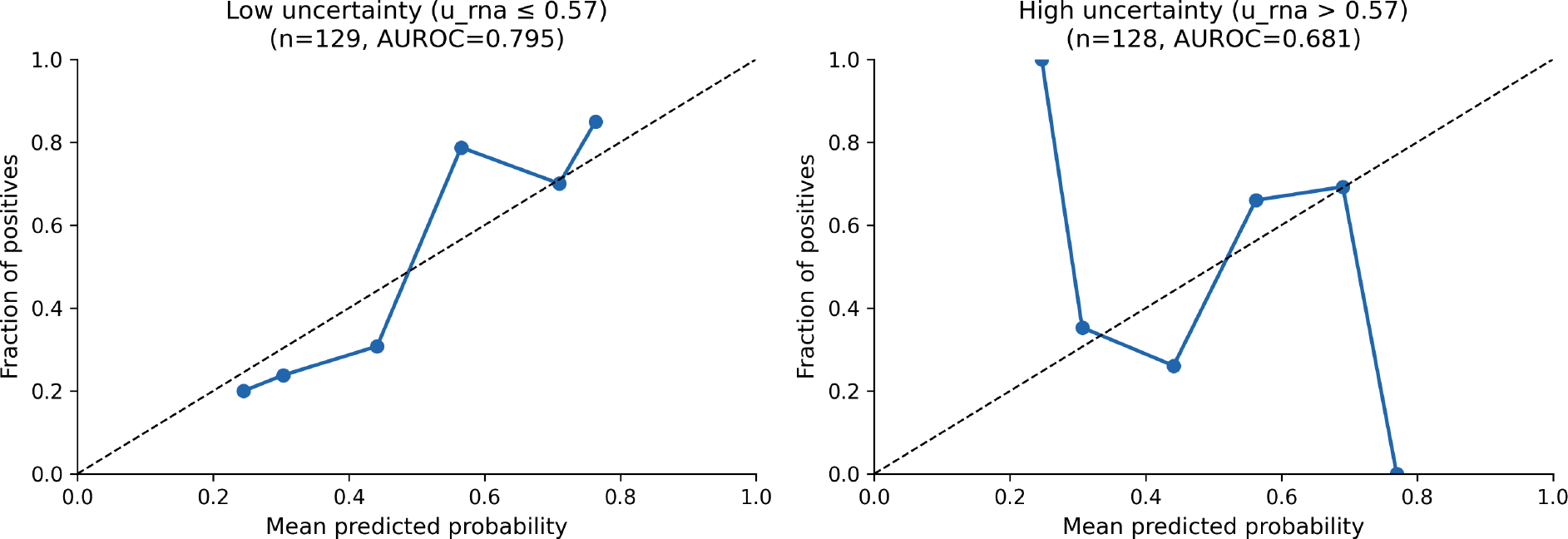
Calibration by uncertainty level. Calibration is stratified by median RNA uncertainty (*u*_RNA_ *≤* 0.57 vs. *>* 0.57). Low-uncertainty patients (AUROC 0.795, *n* = 129) are better discriminated than high-uncertainty patients (AUROC 0.681, *n* = 128), confirming that the uncertainty score is informative. *Note on AUROC aggregation*. The AUROC of 0.772 is the mean of five per-fold AUROCs. The pooled out-of-fold AUROC (combining all 257 classifiable held-out predictions) is 0.730. Both are valid statistics but represent distinct quantities; we report the fold-mean as the primary metric for consistency with C-index and AUPRC.

**Figure 5.**
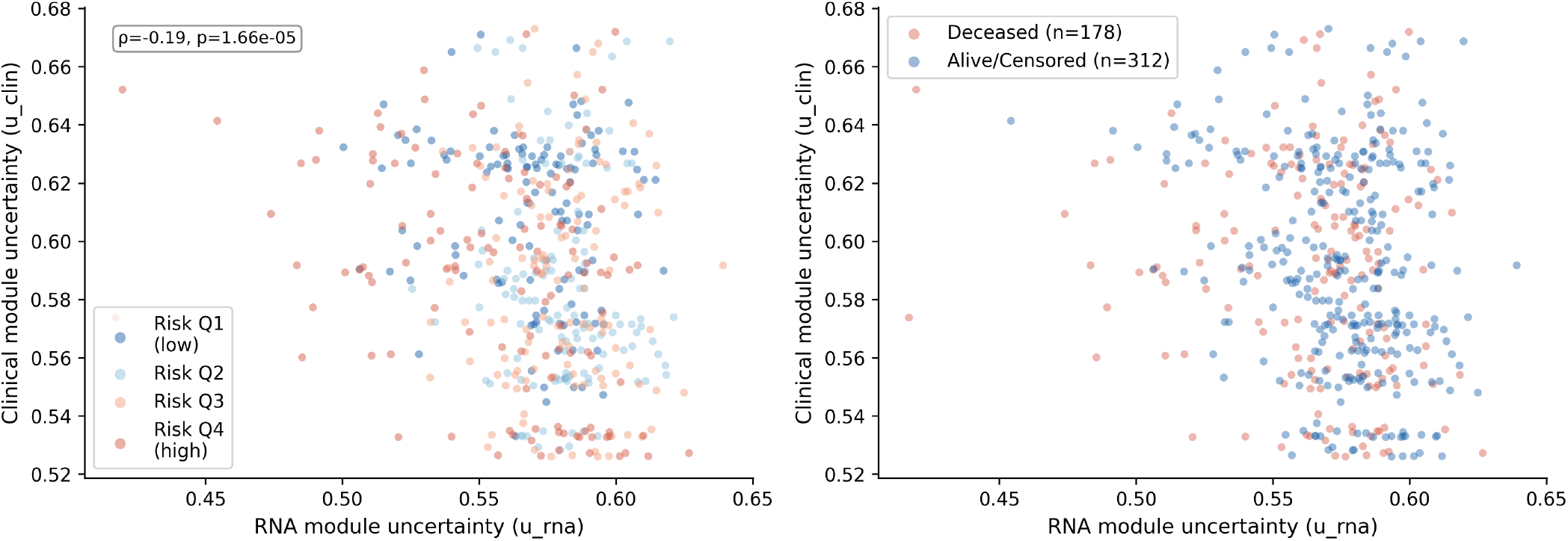
Per-patient epistemic uncertainty (*u*_RNA_ vs. *u*_clin_), pooled across five held-out folds (*n* = 490). Left: coloured by predicted risk quartile. Right: coloured by event status. The negative Spearman correlation (*ρ* = *−*0.19, *p* = 1.66 × 10^*−*5^) shows complementary uncertainty across modalities, consistent with the gate design intent.

### 5.7 Gate Activation by Pathologic Stage

Figure 6 shows mean communication gate activation stratified by pathologic stage. The clinical gate (*g*_clin_) is highest at Stage I (median *≈* 0.510) and shows a monotonic decrease toward Stage IV (median *≈* 0.508), reflecting that at advanced stages, where the clinical stage signal is already maximally informative, the clinical module reduces its receptiveness to transcriptomic messages. The RNA gate (*g*_RNA_) shows a smaller and less monotonic trend, consistent with RNA providing more uniform prognostic signal across stages. This stage-dependent gating behaviour provides mechanistic evidence that the architecture learns meaningful cross-modal dependencies.

**Figure 6.**
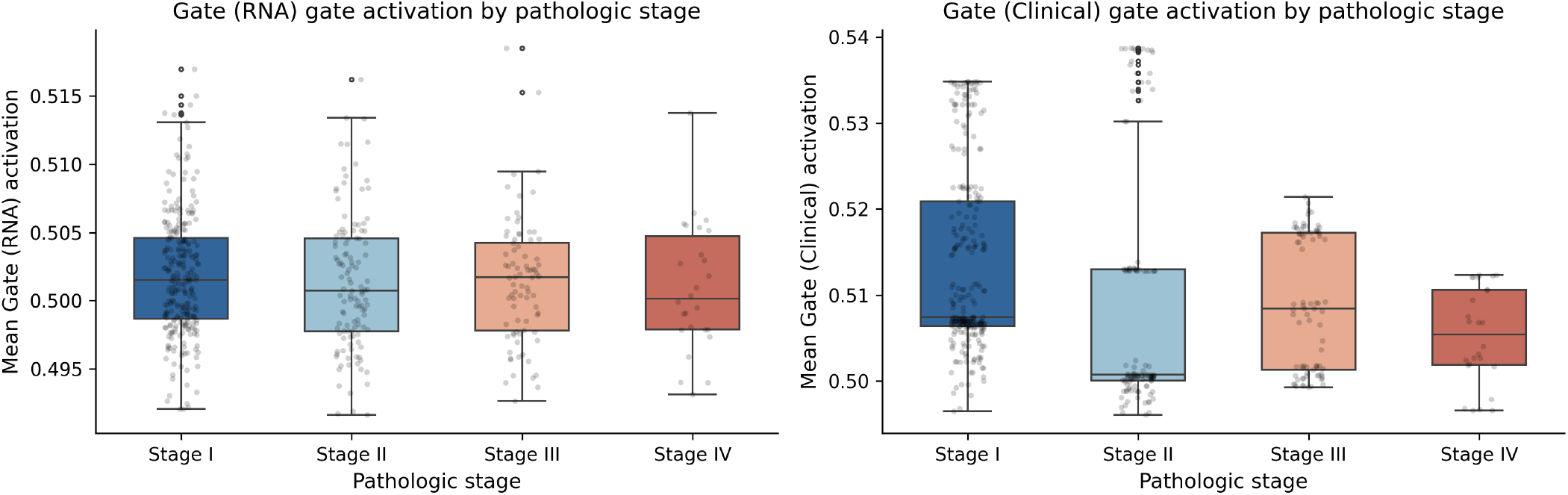
Mean gate activation (RNA gate: left; clinical gate: right) stratified by pathologic stage. The clinical gate decreases from Stage I to Stage IV, indicating reduced receptiveness to transcriptomic messages at stages where clinical features are already strongly prognostic.

### 5.8 Survival-Associated Gene Analysis

Figure 7 shows the top 20 genes by mean absolute Cox coefficient across five training folds (univariate Cox, penaliser 0.1). All top 20 genes carry positive coefficients, indicating risk-increasing expression. The list is dominated by proliferative and metabolic markers: *DKK1* (Wnt signalling antagonist associated with poor prognosis in LUAD [Zhang et al., 2022]), *LDHA* and *GAPDH* (glycolytic enzymes upregulated in proliferative tumours), and cell-cycle regulators *ANLN, HMMR*, and *PLK1*. The biologically motivated markers *EGFR, TP53, SFTPC*, and *AGER* do not appear among the top 20 marginal associations, consistent with their known complex multivariate roles that are attenuated by co-mutation patterns in univariate analysis.

**Figure 7.**
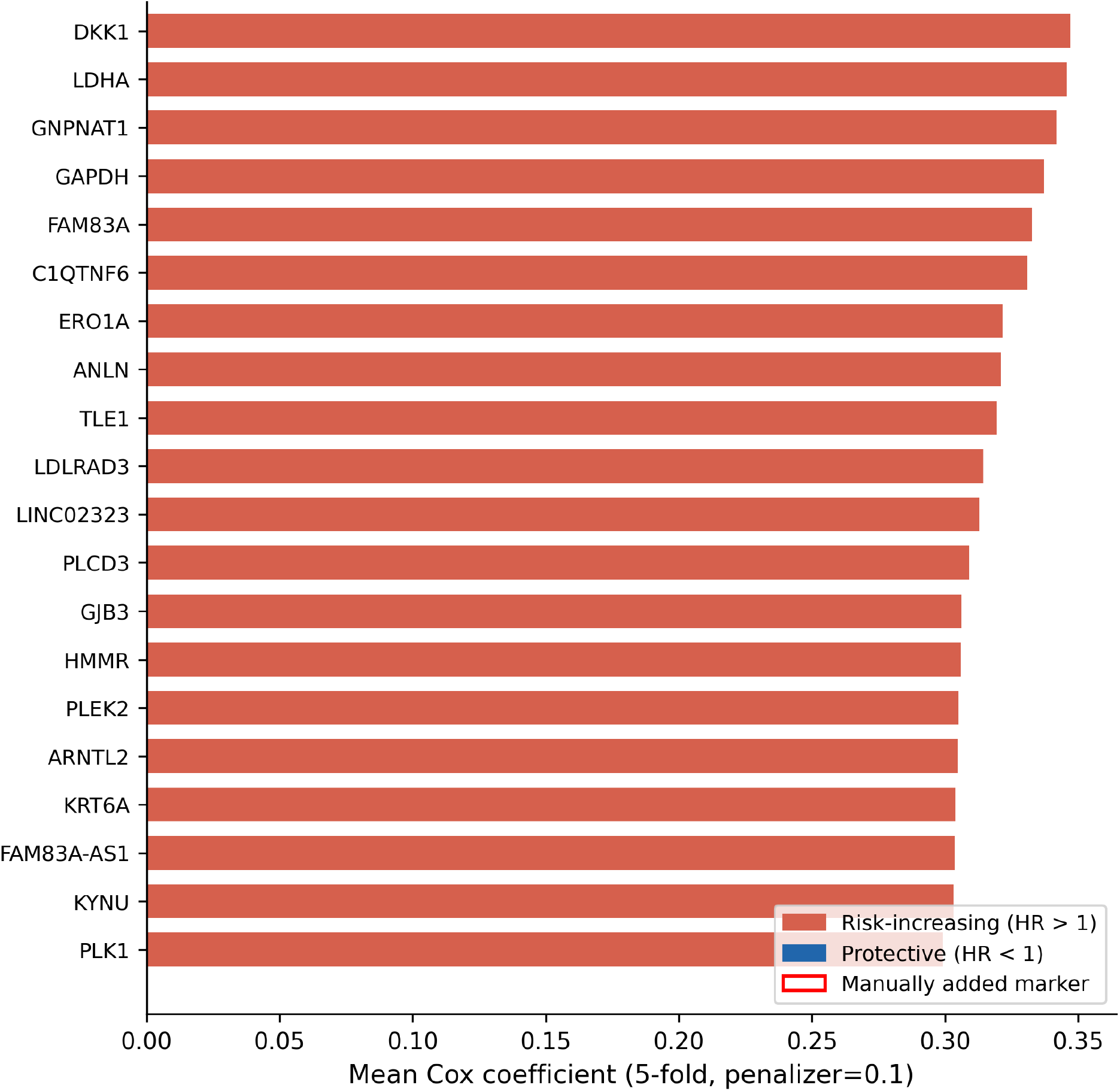
Top 20 survival-associated genes by mean absolute Cox coefficient (5-fold, penaliser = 0.1). All top 20 are risk-increasing (red). Red outline indicates manually added biologically motivated markers. The list is enriched for cell-cycle and glycolytic pathway genes.

## 6 Discussion

### Performance in context

CMRL achieves C-index 0.732 using only RNA-seq and clinical data, surpassing methods with richer modalities (Table 2). The key insight from the ablation is that the concatenation baseline (0.656) performs *worse* than both the RNA-only (0.711) and clinical-only (0.670) unimodal variants. This is a canonical signature of modality collapse: naïve concatenation allowed the high-dimensional RNA representation to suppress the compact clinical signal, actively degrading performance relative to using either modality alone. CMRL resolves this by dynamically regulating information flow through the uncertainty gate, recovering and exceeding the performance of the best unimodal variant by +0.021 C-index.

### Uncertainty interpretation

The negative correlation *ρ* = *−*0.19 between modality uncertainties suggests the two modules are calibrated in a complementary fashion: when the RNA module is uncertain (noisy transcriptomic signal), the clinical module tends to be more certain, and vice versa. The cooperative gate exploits this by routing more information from whichever module is currently confident. The uncertainty-stratified AUROC gap (0.795 vs. 0.681) is the strongest evidence that the uncertainty scores are functioning as intended rather than as uninformative regularisers.

### Calibration

The ECE of 0.122 indicates moderate miscalibration, primarily in the mid-range probability bins (0.35–0.45) where the curve dips below the diagonal. This pattern is consistent with the evidential loss annealing schedule leaving residual overconfidence in lower-probability bins. Post-hoc temperature scaling or isotonic regression should be investigated in future work before clinical deployment.

### Gene analysis

The dominance of glycolytic (*LDHA, GAPDH*) and proliferative (*ANLN, HMMR, PLK1*) markers is consistent with established LUAD biology. The appearance of *DKK1* as the top-ranked gene is notable: DKK1 is a secreted Wnt antagonist whose overexpression drives epithelial-mesenchymal transition and immune evasion in LUAD [Zhang et al., 2022]. These associations are marginal (univariate) and should not be interpreted as causal claims about the prognostic model.

## 7 Conclusion

We presented CMRL, an uncertainty-gated multimodal deep learning framework for LUAD survival prediction combining Evidential Deep Learning, a Variational Information Bottleneck, and an adaptive communication gate. On 490 TCGA-LUAD patients (504-gene RNA-seq, 8-dim clinical features), CMRL achieves C-index 0.732 ± 0.024, AUROC 0.772 ± 0.019, and AUPRC 0.773 ± 0.056 via 5-fold cross-validation. The ablation study reveals that the concatenation baseline (0.656) underperforms both the RNA-only (0.711) and clinical-only (0.670) variants, confirming modality collapse in naïve fusion and validating the CMRL gate design. Calibration analysis (ECE= 0.122) and uncertainty-stratified evaluation (AUROC 0.795 vs. 0.681 for low vs. high uncertainty patients) provide interpretable evidence that the gate mechanism functions as intended. Future work will incorporate whole-slide histopathology as a third module, pursue external validation, and apply post-hoc calibration.

## Acknowledgements

This work was supported by the Research Institute for Medical and Health Sciences (RIMHS), University of Sharjah, Sharjah, United Arab Emirates.

The authors acknowledge the TCGA Research Network for making the LUAD dataset publicly available. Computational experiments were conducted using an NVIDIA Titan RTX GPU.

## Declaration of Competing Interests

The authors declare no competing interests.

## Data Availability

TCGA-LUAD RNA-seq and clinical data are publicly available via the NIH Genomic Data Commons (https://portal.gdc.cancer.gov/). Code and trained model weights will be made available upon acceptance.

## Notes

https://portal.gdc.cancer.gov/

### Competing Interest Statement

The authors have declared no competing interest.

